# ArfGAP1 regulates the endosomal sorting of guidance receptors to promote directed collective cell migration in vivo

**DOI:** 10.1101/2022.09.13.507825

**Authors:** Alison Boutet, Carlos Zeledon, Gregory Emery

## Abstract

Chemotaxis drives diverse migrations important for development and involved in diseases, including cancer progression. Using border cells in the Drosophila egg chamber as a model for collective cell migration, we characterized the role of ArfGAP1 in regulating chemotaxis during this process. We found that ArfGAP1 is required for the maintenance of receptor tyrosine kinases, the guidance receptors, at the plasma membrane. In absence of ArfGAP1, the level of active receptors is reduced at the plasma membrane and increased in late endosomes. Consequently, clusters with impaired ArfGAP1 activity lose directionality. Furthermore, we found that the number and size of late endosomes and lysosomes are increased in the absence of ArfGAP1. Finally, genetic interactions suggest that ArfGAP1 acts on the kinase and GTPase Lrrk to regulate receptor sorting. Overall, our data indicate that ArfGAP1 is required to maintain the homeostasis of the endo-lysosomal pathway to ensure the maintenance of guidance receptors at the plasma membrane and promote chemotaxis.

## INTRODUCTION

Cell migration is a fundamental process that can take various forms. For example, cell migration can be directed or not. One example of directed cell migration is chemotaxis that happens through the binding of chemoattractant by guidance receptors (SenGupta et al., 2021). Furthermore, cells can migrate either individually or collectively, in groups of different sizes (Haeger et al., 2015). Chemotaxis and collective cell migration play fundamental roles during development (Heasman and Ridley, 2008). Unfortunately, they are also mechanisms exploited by cancer cells to form metastasis (Haeger et al., 2015). Understanding how groups of cells sense and respond to chemotaxis gradients is of high importance. Border cell migration in the Drosophila egg chamber is among the most potent models to study collective cell migration and chemotaxis in vivo (Capuana et al., 2020; Montell, 2003; Montell et al., 2012; Peercy and Starz-Gaiano, 2020).

The Drosophila egg chamber is composed of 15 large nurse cells and the oocyte, surrounded by follicle cells. The border cells derive from these somatic cells and form a small cluster of 6-10 cells that perform an antero-posterior, invasive migration in between the nurse cells, toward the oocyte (Montell, 2003; Montell et al., 2012). The oocyte secretes the ligands PVF1, Keren and Spitz. They attract the border cells by activating two receptor tyrosine kinases (RTKs): the epidermal growth factor receptor (EGFR) and PVR, the sole orthologue of the platelet derived growth factor receptor and the vascular endothelial growth factor receptor (Duchek and Rorth, 2001; Duchek et al., 2001; McDonald et al., 2006; McDonald et al., 2003). Upon binding to their ligands, the RTKs recruit the Rac guanine exchange factor (GEF) Vav to activate Rac and form protrusions in 1 or 2 cells at the leading edge of the cell cluster (Fernandez-Espartero et al., 2013). Rac activity and protrusion formation seem to be actively repressed in the other cells of the cluster through a mechanism involving the actin and plasma membrane binding protein Moesin, the cell-cell adhesion protein DE-cadherin and Myosin II-mediated contractility (Mishra et al., 2019; Plutoni et al., 2019; Ramel et al., 2013; Roberto and Emery, 2022; Wang et al., 2020).

In previous work, we and others have shown that vesicular trafficking plays an important role in regulating border cell migration (Assaker et al., 2010; Cobreros-Reguera et al., 2010; Janssens et al., 2010; Jekely et al., 2005; Laflamme et al., 2012; Ramel et al., 2013; Wan et al., 2013; Zeledon et al., 2019). In particular, endocytosis and recycling were shown to regulate the level and the distribution of active RTKs at the plasma membrane and hence to be required for directed migration (Assaker et al., 2010; Janssens et al., 2010; Jekely et al., 2005; Laflamme et al., 2012; Wan et al., 2013). In mammals, the trafficking of RTKs was shown to be regulated by several means (reviewed in (Caldieri et al., 2018)). For example, the ubiquitination of the EGFR by the E3-ligase Cbl changes its trafficking in the endocytic pathway. Indeed, while ubiquitination is dispensable for the internalisation of RTKs from the plasma membrane, it serves as a sorting signal in endosomes, where ubiquitinated RTKs interact with the ESCRT-0 complex, composed of Hrs and Stam. This leads to the sorting of RTKs into internal vesicles of multivesicular bodies and, subsequently, to transport to late endosomes and lysosomes, where they are degraded. On the contrary, non-ubiquitinated receptors can be recycled to the plasma membrane. Accordingly, border cells mutant for *cbl* have increased levels of RTKs at the plasma membrane suggesting either that less receptors are endocytosed or that they are recycled instead of being degraded. Furthermore, Hrs and Stam loss-of-functions in Drosophila lead to an accumulation of active RTKs in endosomes (Jekely et al., 2005). Overall, this suggests that a mechanism similar to the regulation of EGFR in mammals is at play to eliminate ubiquitinated receptors in flies. In addition, the recycling of RTKs was also observed in border cells, to maintain active RTKs at the cortex. Accordingly, Rab GTPases regulating entry and recycling from endosomes (Rab4, Rab5 and Rab11), their regulators (the GEF Sprint and the GTPase activating proteins (GAPs) Evi5 and Rn-Tre) and their effectors (the exocyst complex) have also been involved in border cell migration (Assaker et al., 2010; Janssens et al., 2010; Laflamme et al., 2012; Wan et al., 2013). However, the exact mechanism by which RTKs are sorted toward the different endocytic compartments during border cell migration is still unclear to our knowledge.

To increase our understanding of the regulation of vesicular trafficking in border cells, we have previously performed a candidate RNAi screen (Zeledon et al., 2019). This screen was directed against Arf, Arf-like GTPases and their regulators, as Arf GTPases are known to be involved in the sorting of cargoes into vesicles (Adarska et al., 2021). Despites several candidates inducing pleiotropic effects, as expected since Arf GTPases are required to form vesicles in the biosynthetic pathway (Jackson and Bouvet, 2014; Rodrigues and Harris, 2019), we found that a few candidates induced specific phenotypes. For example, we found that the GAP Drongo was required to promote contractility at the onset of border cell migration (Zeledon et al., 2019). In the present study, we focus on ArfGAP1. ArfGAP1 was originally shown to act as a GAP for Arf1 (Cukierman et al., 1995). As Arf1 is necessary for the formation of COPI-coated vesicles, ArfGAP1 was shown to be an essential regulator of their formation (Lee et al., 2005; Liu et al., 2005). In Drosophila, the ArfGAP1 activity toward Arf1 was involved in eye pigmentation and rhabdomere biogenesis (Raghu et al., 2009; Rodriguez-Fernandez and Dell’Angelica, 2015), in blood cell homeostasis (Khadilkar et al., 2014) and in epithelial tube expansion in the trachea (Armbruster and Luschnig, 2012). Interestingly, the mammalian orthologue of ArfGAP1 was also shown to act as a GAP for Lrrk2 (Stafa et al., 2012; Xiong et al., 2012). Lrrk proteins have both a GTPase and kinase activity (Civiero and Bubacco, 2012) and Lrrk2 is an important factor involved in the development of Parkinson Disease (Usmani et al., 2021). Here, we show that ArfGAP1 is required for the chemotaxis of border cells by maintaining active RTKs at the cell surface. This is achieved by controlling the homeostasis of the endo-lysosomal pathway possibly through the regulation of Lrrk, the sole Lrrk1 and Lrrk2 orthologue in flies.

## RESULTS

### The GAP activity of ArfGAP1 is required for border cell migration

In previous work (Zeledon et al., 2019), we have performed an RNAi screen to identify new regulators of collective cell migration among Arf and Arf-like GTPases, their GAPs and their GEFs. In this screen, we used the Gal4/UAS system to express the control dsRNAs or dsRNAs targeting candidates specifically in border cells with the *c306*-Gal4 driver. ArfGAP1 caught our attention as its depletion impaired border cell migration, but did not induce pleiotropic effects and did not disrupt the Golgi apparatus (Fig.1A and B, Fig.S1A-C and (Zeledon et al., 2019)). To validate that the phenotype was not due to off-targets, we overexpressed a mCherry-tagged version of ArfGAP1 in depleted clusters. Although this construct is still targeted by the RNAi line, we observed a complete rescue of border cell migration (Fig.1A and B).

**Figure 1:**
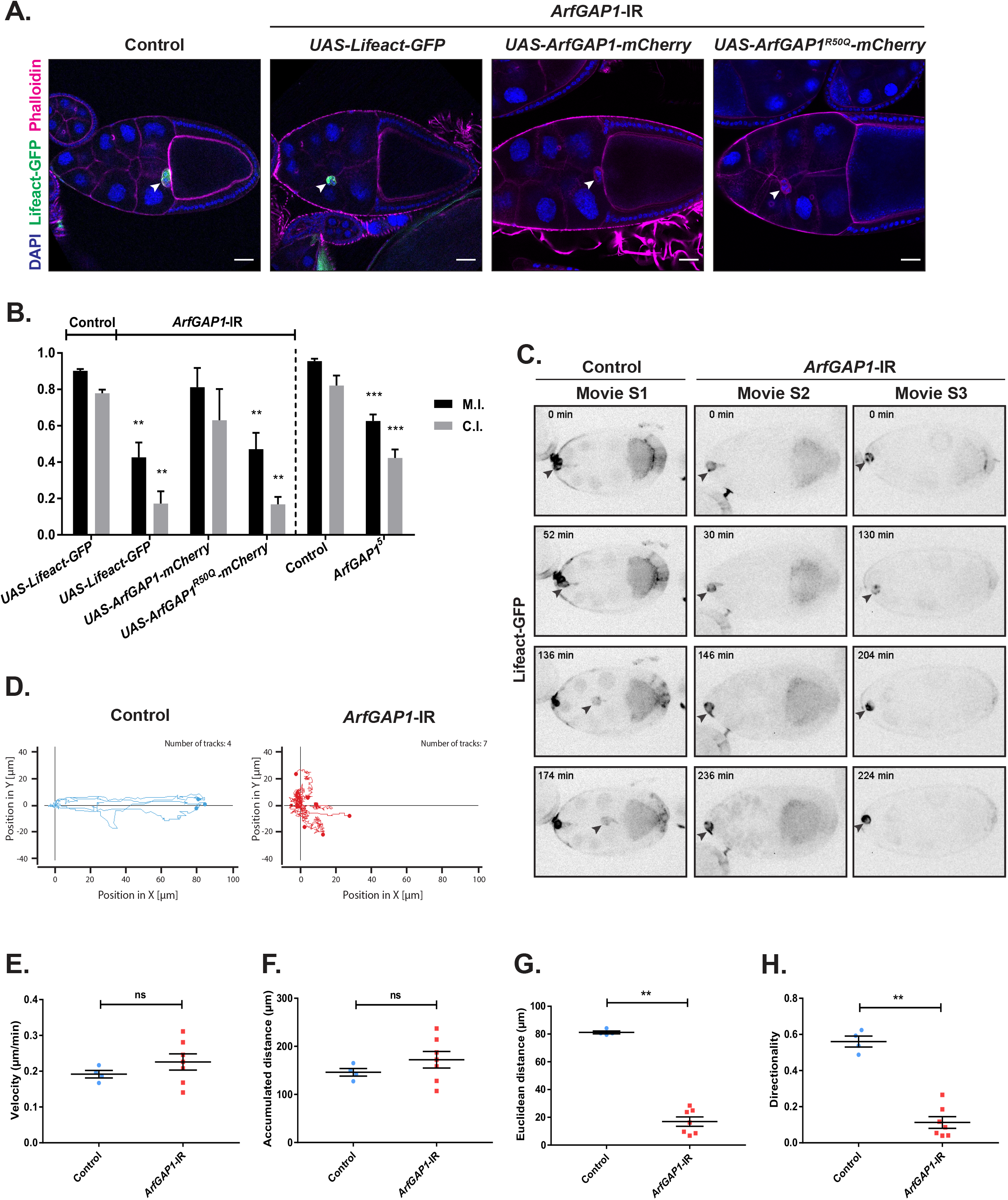
ArfGAP1 is required for the directional migration of border cells (A) Representative images of a control stage 10 egg chamber and stage 10 egg chambers expressing the ArfGAP1 RNA interference (*ArfGAP1-*IR) with the indicated constructs. Arrowheads indicate the localization of the border cell cluster. Scale bar: 30 μm. (B) Migration Indexes (M.I.) and Completion Indexes (C.I.) (see materials and methods) for the different genotypes indicated. Error bars are SEM (N=3 experiments; **: p<0.01, ***: p<0.001, ****: p<0.0001, One-way ANOVA test). (C) Representative images from time-lapse recordings of the migration of border cells expressing Lifeact-GFP in (left, Movie S1) control egg chamber or (middle, right, Movie S2 and S3) after ArfGAP1 depletion. Arrowheads indicate the localization of the border cell cluster. (D) Traces from time-lapse recordings of border cell migration in control egg chambers (left) or after depletion of ArfGAP1 (right). (E-H) Mean velocity, mean accumulated distance, mean Euclidian distance, and mean directionality for control and ArfGAP1 depleted clusters. Error bars are SEM (n.s.: not significant (p>0.05), **: p<0.01, Mann-Whitney test).

To confirm that the loss of ArfGAP1 blocks border cell migration, we generated a null mutant by using CRISPR/Cas9 (Zirin et al., 2020). We isolated different depletions. Among these, the allele *ArfGAP1*^*5*^ has a frameshift at amino acid 37 leading to a stop codon at position 75. Consequently, *ArfGAP1*^*5*^ has lost the catalytic arginine at position 50 and should be non-functional. *ArfGAP1*^*5*^ is homozygous viable and induces a delayed border cell migration that was similar to the depletion by RNAi (Fig.1B). To determine if *ArfGAP1*^*5*^ acts as genetic nulls, we crossed it to a deficiency line covering *ArfGAP1* (Df(3L)BSC730). The phenotype observed in *ArfGAP1*^*5*^*/*Df(3L)BSC730 was identical to homozygous *ArfGAP1*^*5*^, indicating that *ArfGAP*^*5*^ is a null allele.

To determine if the GAP activity of ArfGAP1 is required for border cell migration, we generated a GAP inactive form of ArfGAP1 (ArfGAP1^R50Q^) and expressed it in ArfGAP1 depleted clusters. We found that the expression of the catalytic inactive ArfGAP1 was unable to rescue ArfGAP1 depletion, showing that the GAP activity is necessary for ArfGAP1’s function in border cells (Fig.1A and B).

### ArfGAP1 is necessary to maintain the directionality of border cell migration

To gain information, we compared time-lapse recordings of control clusters to clusters depleted for ArfGAP1. Contrary to control clusters (Movie S1, Fig.1C)) that perform a direct migration toward the oocyte, we observed two types of behaviors after the depletion of ArfGAP1: 1) the cluster seemed active but was not progressing, possibly undergoing a shuffling and tumbling behavior as control clusters undergo when they probe for directionality (Bianco et al., 2007) (Movie S2, Fig.1C), or 2) the cluster started migrating but then lost directionality (Movie S3, Fig.1C). In the later case, we always saw a normal number of protrusions polarized at the front of migration and cohesive clusters. Accordingly, we found that the active, phosphorylated form of Moesin that is required to coordinate the cells of the cluster (Plutoni et al., 2019; Ramel et al., 2013) is also normally distributed at the periphery of the cluster after the depletion of ArfGAP1 (Fig.S1D). We quantified time-lapse recordings of clusters depleted of ArfGAP1 and found that the mean velocity of depleted clusters was 0.225838µm/min, similar to control cluster (0.191683µm/min) but the directionality and the total displacement for the course of the recording were dramatically reduced (Fig.1D-H).

Loss of directionality can be due to abnormal polarization of the cluster. Hence, we monitored the distribution of the polarity proteins Discs Large and Bazooka (Par3) that are required for the spatial organization of the cluster (Li et al., 2009; Pinheiro and Montell, 2004; Wang et al., 2018) and found that they were unaffected (Fig.S1E-F). The migration phenotype we observed was similar to the loss-of-function of DE-cadherin which is necessary for the border cell - nurse cell interaction, to maintain the cohesion in the cluster and to ensure directional migration (Cai et al., 2014; Niewiadomska et al., 1999). We found that DE-cadherin is normally distributed in ArfGAP1 depleted clusters (Fig.S1G). Similarly, another adhesion protein, Fasciclin III (Han et al., 2000) was also unaffected by the depletion of ArfGAP1 (Fig.S1H). Overall, this indicates that most of the determinants required for border cell migration are unaffected by the depletion of ArfGAP1. Hence, we decided to further investigate the exact molecular mechanism regulated by ArfGAP1 in border cells.

Since directionality is lost, but polarity is normal, we hypothesized that the capacity of ArfGAP1 depleted clusters to sense the gradient of RTK ligands guiding border cell migration is impaired. Indeed, clusters having impaired RTK activity are unable to sense the ligand gradient (Prasad and Montell, 2007), and clusters migrating in egg chambers expressing uniform levels of PVF1 migrate abnormally, displaying increased shuffling behavior (Bianco et al., 2007).

### Depletion of ArfGAP1 leads to an accumulation of active RTKs in the endo-lysosomal pathway

To test the hypothesis that ArfGAP1 regulates RTKs, we monitored the distribution of active RTKs at the onset of migration with the monoclonal antibody 4G10 that recognizes phosphorylated Tyrosine (pTyr) (Assaker et al., 2010; Jekely et al., 2005; Laflamme et al., 2012). While in control conditions active RTKs are predominantly found at the cortex, after depletion of ArfGAP1 or in homozygous *ArfGAP1*^*5*^ mutant egg chambers, the pTyr signal accumulated in intracellular, vesicular-like structures (Fig.2A-B). Concomitantly, we observed a decrease of active RTKs at the plasma membrane at the migration front (Fig.2C). To test if this redistribution is due to the loss of the GAP activity of ArfGAP1, we performed rescue experiments. We found that the expression of wildtype ArfGAP1 in ArfGAP1 depleted clusters restores the normal localization of active RTKs that are now excluded from endosomal structures and found at the plasma membrane. On the contrary, the expression of the catalytic inactive form of ArfGAP1 did not rescue the phenotype indicating that the catalytic activity of ArfGAP1 is important to maintain active RTKs at the plasma membrane (Fig.2A-C).

**Figure 2:**
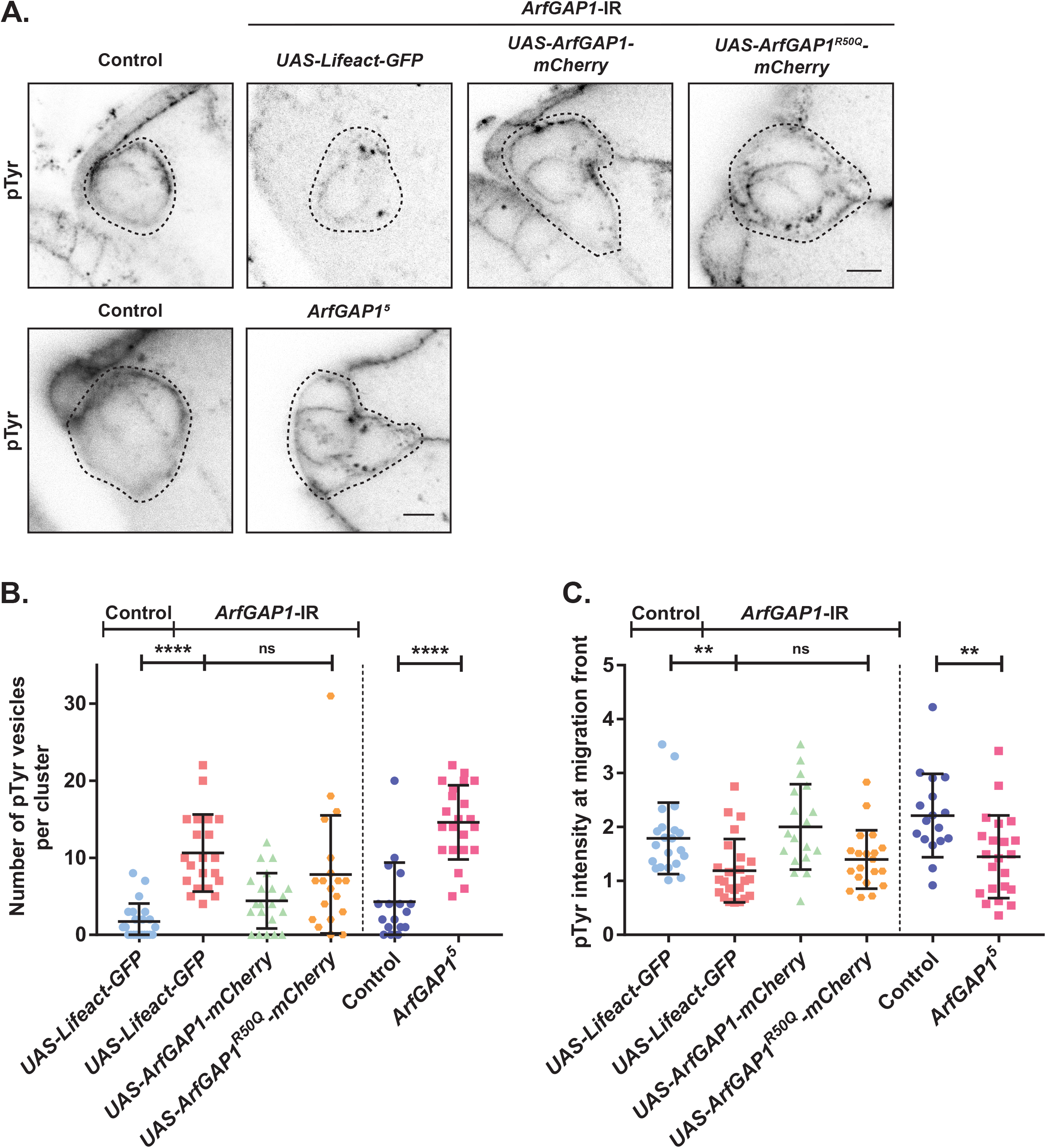
ArfGAP1 depletion mis-localizes active RTKs in vesicular structures (A) Representative images of border cell clusters of the indicated genotype, labeled with the anti-pTyr antibody 4G10. Scale bar: 5 µm. Clusters are delineated by a dotted line. (B) Quantification of the number of pTyr positive vesicles per cluster in the indicated conditions. Error bars are SEM (6≤n≤12 clusters; N=3 experiments; n.s.: not significant (p>0.05), ****: p<0.0001, Kruskal-Wallis and Mann-Whitney test). (C) Quantification of the mean plasma membrane intensity of the pTyr signal at the migration front (see material and methods) in the indicated conditions. Error bars are SEM (6≤n≤12 clusters; N=3 experiments; n.s.: not significant (p>0.05), **: p<0.01, Kruskal-Wallis and Mann-Whitney test).

Next, we investigated in which compartment active RTKs were trapped after ArfGAP1 depletion by costaining border cells with pTyr and markers of various endocytic compartments. First, we observed that active RTKs co-localized with large structures labelled with the endosomal marker GFP-myc2xFYVE (Wucherpfennig et al., 2003) (Fig.3A). More precisely, active RTKs were found inside structures surrounded by FYVE. This suggested that active RTKs were accumulating in multivesicular bodies or in late endosomes. Accordingly, we found that pTyr did not co-localize with markers of the early endosome (GFP-Rab5, (Wucherpfennig et al., 2003)) nor with GFP-Sec15, a marker of the recycling endosomes (Jafar-Nejad et al., 2005) (Fig.3B-C). However, we found that pTyr accumulated inside GFP-Rab7-positive structures, possibly in intralumenal vesicles (Fig.3D). Co-labelling with the lysosomal marker LysoTracker showed that the pTyr signal was not found inside lysosomes, but frequently juxtaposed to lysosomal structures (Fig.3E). We hypothesize that RTKs are degraded when entering lysosomes and thus cannot be detected in them. We found a very similar distribution of active RTKs in *ArfGAP1*^*5*^ homozygous egg chambers (Fig.3F). Altogether, these data suggest that active RTKs are overly targeted in the degradative pathway in the absence of ArfGAP1.

**Figure 3:**
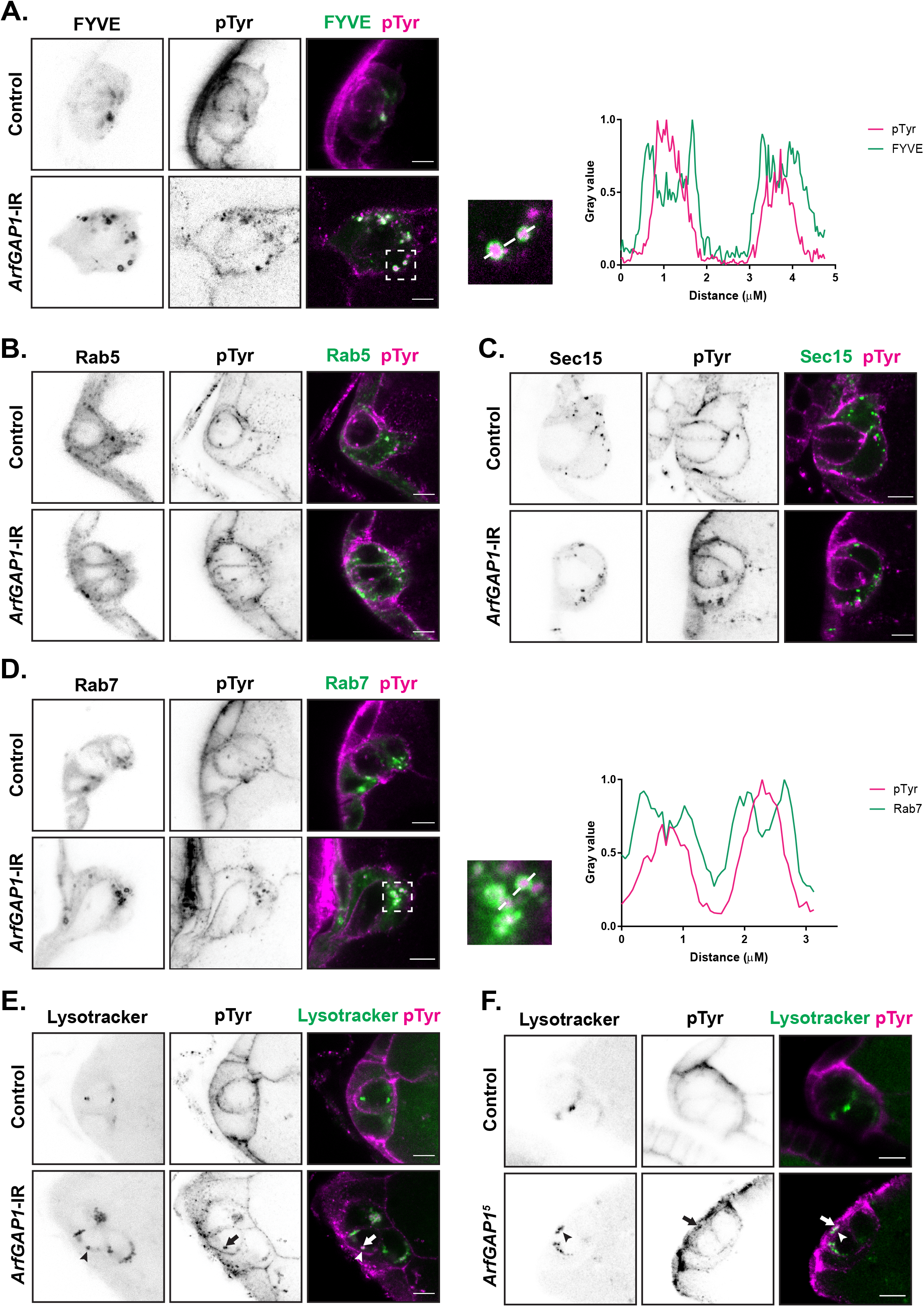
Active RTKs (pTyr) accumulate in late endosomes after the depletion of ArfGAP1. (A) Co-labelling of pTyr by immunofluorescence and endosomes (marked with GFP-myc2xFYVE) in control and ArfGAP1 depleted clusters. On the right, the high magnification image and the quantification of fluorescence along a line that span over two puncta reveal that the pTyr signal is inside FYVE-positive vesicles. Scale bar: 5 µm. (B) Co-labelling of pTyr and early endosomes (GFP-Rab5) and (C) co-labelling of pTyr with recycling vesicles (GFP-Sec15) show no obvious overlap, even after the depletion of ArfGAP1. Scale bar: 5 µm. (D) Co-labelling of pTyr with the late endosomal marker GFP-Rab7. On the right, the high magnification image and the quantification of fluorescence by “line-scan” reveal that the pTyr signal is inside Rab7-positive vesicles in ArfGAP1-depleted clusters. Scale bar: 5 µm. Co-labelling of pTyr with LysoTracker in control clusters, after depletion of ArfGAP1 (E) or in homozygous *ArfGAP1*^*5*^ egg chambers (F). Scale bar: 5 µm. Arrowheads and arrows indicate the localization of Lysotracker and pTyr vesicles respectively that are juxtaposed.

### ArfGAP1 is required for the homeostasis of the degradative pathway

Furthermore, we observed that the apparent GFP-Rab7 signal was increased in clusters depleted for ArfGAP1 (Fig.3D). We confirmed this observation by quantifying the mean intensity of the GFP-Rab7 signal at the onset of migration (Fig.4A). Correlating with our observation that active RTKs accumulate in late endosomes, we also found a significant increase of the pTyr signal inside Rab7 vesicles when ArfGAP1 is depleted (Fig.4B). To document a possible increase of the endo-lysosomal degradative pathway, we quantified the size of late endosomes and lysosomes as marked by Rab7 and LysoTracker, respectively and we found a significant increase after ArfGAP1 depletion (Fig.4C-D). Moreover, the number of Lysotracker vesicles increased significantly in that condition (Fig.4E). To verify that this was indeed due to the loss of ArfGAP1, we measured the size and the number of lysosomes in homozygous *ArfGAP1*^*5*^ clusters. We also observed a similar increase of the number and the size of lysosomes in *ArfGAP1*^*5*^ mutants (Fig.4D-E). These data show that ArfGAP1 is required for the homeostasis of the degradative pathway.

**Figure 4:**
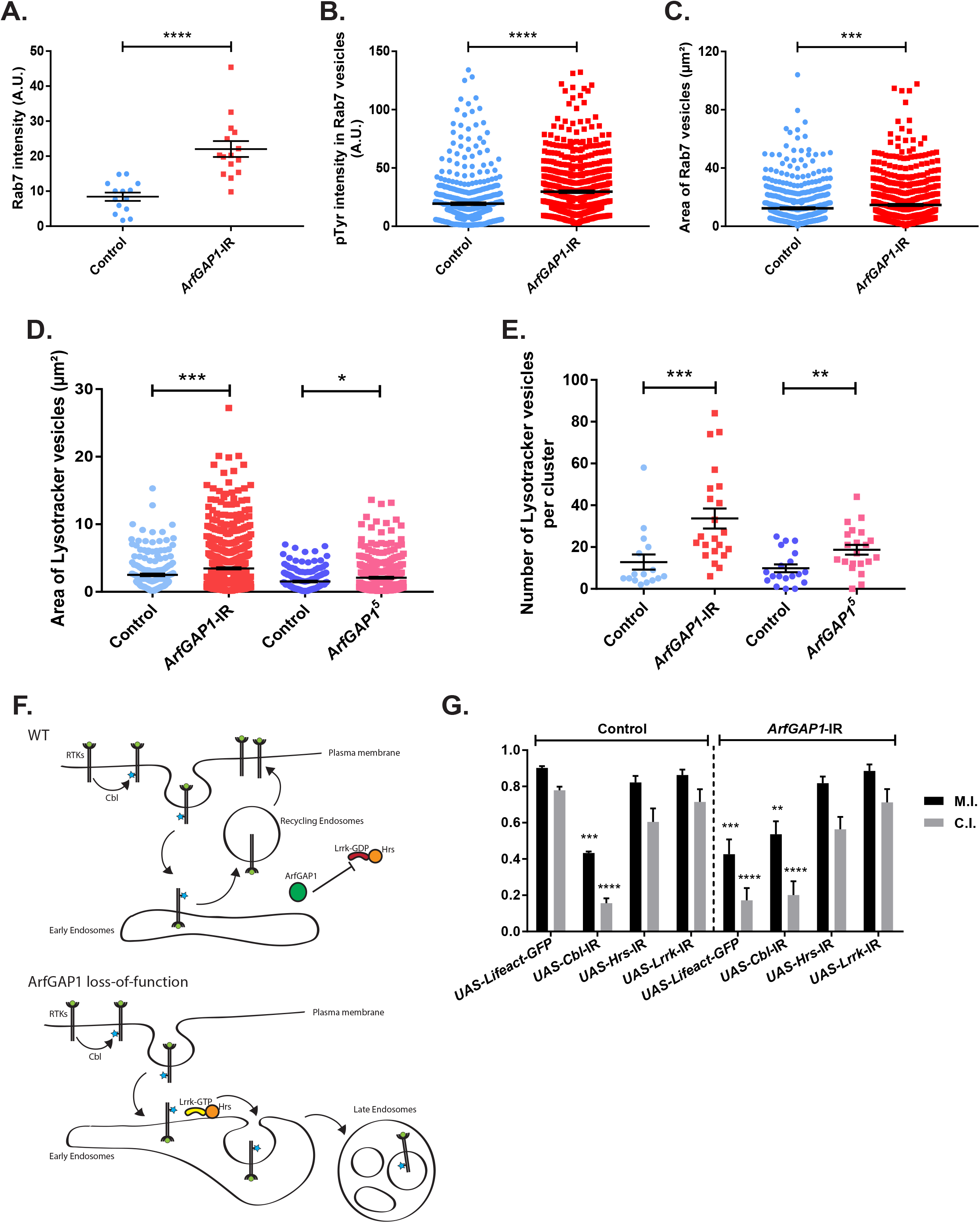
ArfGAP1 regulates the homeostasis of the endo-lysosomal degradative pathway and interact genetically with Lrrk and Hrs. (A) Quantification of the GFP-Rab7 intensity in control cluster and after depletion of ArfGAP1. Error bars are SEM (4≤n≤8 clusters; N=3 experiments; ****: p<0.0001, Mann-Whitney test). (B) Intensity of the pTyr signal within Rab7-positive structures in control clusters and after depletion of ArfGAP1. Error bars are SEM (N=3 experiments; ****: p<0.0001, Mann-Whitney test). (C) Area covered by Rab7-positive vesicles in control clusters and after depletion of ArfGAP1. Error bars are SEM (N=3 experiments; ***: p<0.001, Mann-Whitney test). (D) Area covered by LysoTracker positive vesicles in control clusters, after depletion of ArfGAP1 and in homozygous *ArfGAP1*^*5*^ egg chambers. Error bars are SEM (N=3 experiments; *: p<0.05, ***: p<0.001, Kruskal-Wallis test). (E) Number of LysoTracker positive vesicles in control clusters, after depletion of ArfGAP1 and in homozygous *ArfGAP1*^*5*^ egg chambers. Error bars are SEM (4≤n≤10 clusters; N=3 experiments; **: p<0.01, ***: p<0.001, Kruskal-Wallis test). (F) Proposed model depicting how ArfGAP1 regulates the trafficking of RTKs. (G) Migration and completion indexes of border cell migration in the indicated conditions. Error bars are SEM (N=3 experiments; **: p<0.01, ***: p<0.001, ****: p<0.0001, One-way ANOVA test).

### The ArfGAP1 phenotype is rescued by the co-depletion of Lrrk or Hrs

Our findings suggest that ArfGAP1 inhibits the degradative pathway and the degradation of RTKs. In ArfGAP1 loss of function, RTKs would thus be overly degraded (Fig.4F). To determine the molecular mechanism regulating the sorting of RTK that is regulated by ArfGAP1, we used genetic interactions. In control conditions, a pool of RTKs is ubiquitinated by the E3 ubiquitin ligase Cbl, which promotes their endocytosis and degradation (Jekely et al., 2005). We hypothesize that in ArfGAP1 depleted cluster, most of the ubiquitinated RTKs are now targeted into the degradative pathway. If our hypothesis is accurate, depleting Cbl may rescue the ArfGAP1 phenotype by reducing the endocytosis and degradation of RTKs. We found that Cbl co-depletion does not rescue border cell migration in ArfGAP1 depleted clusters (Fig.4G). As depletion of Cbl induces a migration phenotype *per se*, the absence of genetic interaction between ArfGAP1 and Cbl does not allow us to conclude that ArfGAP1 and Cbl act independently. However, this result suggests that the ArfGAP1 migration phenotype is not entirely dependent on the ubiquitination of RTKs.

Next, to gain insight into the mechanism by which ArfGAP1 regulates the localization of active RTKs, we searched for potential substrates of ArfGAP1 in the literature. ArfGAP1 was shown to act as a GAP on Arf1 and Lrrk2 (Cukierman et al., 1995; Stafa et al., 2012; Xiong et al., 2012). As ArfGAP1 depletion does not affect the Golgi apparatus, to the difference of Arf1 depletion (Fig.S1A-C and (Zeledon et al., 2019)), we thought that it is unlikely that ArfGAP1 acts on Arf1 to regulate border cell migration. On the other hand, Lrrk1 was shown, in mammals, to promote the transport of EGFR from early to late endosomes by binding to Hrs and Stam, two components of the ESCRT-0 complex (Hanafusa et al., 2011; Ishikawa et al., 2012). In Drosophila, there is a single Lrrk1 and Lrrk2 orthologue (Lrrk). Hence, it is possible that in the absence of ArfGAP1, Lrrk is overactivated and promotes the trafficking of RTKs into late endosomes and lysosomes through the recruitment of the ESCRT-0 complex (Fig.4F). To test this, we used genetic interactions. We rationalized that if the depletion of ArfGAP1 increases Lrrk activity, the co-depletion of Lrrk should restore migration. Accordingly, we found that the ArfGAP1 phenotype is rescued by the co-depletion of Lrrk (Fig.4G). Similarly, we co-depleted Hrs and ArfGAP1 and observed a rescue of the migration phenotype (Fig.4G). Overall, our data suggests that ArfGAP1 acts on Lrrk to inhibit the ESCRT-0-mediated sorting of RTKs into the endo-lysosomal degradative pathway and their subsequent degradation.

## DISCUSSION

Chemotaxis plays fundamental roles during development and in diseases, such as cancer progression. During border cell migration, vesicular trafficking was shown by us and others to spatially control the distribution of the receptors responsible for chemotaxis. A recent screen performed in our laboratory identified new regulators of border cell migration including two Arf GAPs: Drongo and ArfGAP1 (Zeledon et al., 2019).

Here, we focused on ArfGAP1, which has been shown to act as a GAP on Arf1 (Cukierman et al., 1995) and on Lrrk2 (Stafa et al., 2012; Xiong et al., 2012). Our findings suggest that ArfGAP1 acts on Lrrk in border cells. Lrrk proteins are composed of both a kinase and a GTPase domain. Mammals have two Lrrks (Lrrk1 and Lrrk2) that were shown to phosphorylate GTPases of the Rab family to modulate their interactions with specific effectors (Malik et al., 2021). For example, Lrrk1, Lrrk2 and Drosophila Lrrk were shown to regulate Rab7 (Dodson et al., 2012; Gomez-Suaga et al., 2014; Hanafusa et al., 2019; Malik et al., 2021). Phosphorylation of Rab7 by Lrrk1 modulates its bindings to its effector RILP, that links Rab7 to dynein and regulates the positioning of late endosomes within the cell (Hanafusa et al., 2019). Accordingly, the expression of a Lrrk gain-of-function in *Drosophila* follicle cells inhibits the perinuclear accumulation of Rab7-positive puncta, and increases the size of late endosomes and lysosomes in a Rab7-dependent mechanism (Dodson et al., 2012). Hence, it is appealing to hypothesize that Rab7 is a key target of Lrrk in border cells. This would need to be tested in detail.

Interestingly, mammalian Lrrk1 was shown to regulate the transport of EGFR from early to late endosomes (Hanafusa et al., 2011; Hanafusa et al., 2019; Ishikawa et al., 2012). In particular, Lrrk1 can interact with EGFR and the ESCRT-0 constituents Hrs and Stam in early endosomes (Hanafusa et al., 2011). Our work suggests that in border cells, ArfGAP1 constrains the activity of Lrrk to inhibit the ESCRT-0-mediated sorting of RTKs into multivesicular bodies and the degradative pathway. Combined with these previous studies, our work suggests that ArfGAP1 controls the endosomal sorting of RTKs through its GAP activity toward Lrrk that phosphorylates Rab7 and possibly other Rab proteins, and/or recruits Hrs and Stam. Our model also suggests that the recycling of RTKs is promoted in the presence of ArfGAP1 (Fig.4F). In the context of border cell migration, ArfGAP1 plays thus an important role in maintaining active RTKs at the plasma membrane, and this ensures the proper guidance of border cells. It is interesting to speculate that ArfGAP1 and Lrrk are regulators of the trafficking of RTKs in various contexts. As RTKs are required for various processes during development, tissue homeostasis and in the development of cancers, understanding the mechanism of action of ArfGAP1 and Lrrk on RTKs is important.

An unexpected finding was the increase in late endosomes and lysosomes in ArfGAP1 loss of function conditions. This suggests that normal levels of ArfGAP1 are required to maintain the entire degradative pathway to its basal level. It would be important to address in future work if this role of ArfGAP1 is cell specific to border cells or acts as a general mechanism for lysosomal homeostasis. Furthermore, it would be interesting to determine if ArfGAP1 affects the protein levels of other transmembrane. In border cells, we observed that it does not reduce DE-cadherin levels at the cortex (Fig.S1G), demonstrating that some cargoes are still normally recycled. Furthermore, the recycling endosome compartment seems morphologically normal in ArfGAP1 loss of function (Fig.3C and S1I). Overall, this suggests that the stimulation of the degradative pathway in ArfGAP1 loss of function is not due to an impairment of the recycling pathway. Indeed, if this would be the case, we would anticipate an atrophy of the recycling pathway.

Positive feedback loops have long been proposed as a mechanism to transform a shallow extracellular gradient into a strong intracellular gradient, including during border cell migration (Jekely et al., 2005; Wan et al., 2013). Indeed, a positive feedback loop might be an efficient way of transforming a signalling cue into a robust, directed response. As we found that ArfGAP1 regulates RTKs maintenance at the plasma membrane, it would be interesting to determine if it is part of a feedback loop that would be more active in the leader cell than in follower cells, and hence would reinforce the response to extracellular signals.

## MATERIAL AND METHODS

### Drosophila genetics

Crosses were performed at 25°C and flies were incubated at 29°C for 48h before dissection. Knockdown and rescue experiments were performed with the c306-GAL4 driver. This driver expresses specifically in border cells. As we study migration in the ovary, the experiments were conducted in females. Transgenic flies were generated by BestGene Inc. The *ArfGAP1*^*5*^ allele was generated by CRISPR using the TRiP-CRISPR Knockout stocks: nos-Cas9 (#78781) and sgRNA ArfGAP1 (#81490) obtained from the Bloomington Stock Collection. To identify the sequence alterations, we isolated DNA using the QIAGEN DNeasy Blood & Tissue Kit (QIAGEN, Hilden, Germany), amplified it by RT-PCR and sequenced the targeted site using 5’-GCGCCAGCGTCAGCAAAAGTTTCATTAGC-3’ as forward primer and 5’CCTCCTTGAGATCCCAGCTCTTGCCC3’ as reverse primer.

The following stocks were obtained from the Bloomington *Drosophila* Stock Center: *c306*-GAL4 (#3743), UAS-Lifeact-GFP (#35544), UAS-RNAi mCherry (#35785), UAS-RNAi Lrrk (#39019). The fly lines: UAS-RNAi ArfGAP1 (#26460), UAS-RNAi Cbl (#330142) and UAS-RNAi Hrs (#330597) were obtained from the Vienna Drosophila RNAi Center (Dietzl et al., 2007). UAS-GFP-myc2xFYVE/TM3, UAS-GFP-Rab5/TM3, UAS-GFP-Rab7/CyO were obtained from the group of M. Gonzalez-Gaitan (Department of Biochemistry, Université de Genève). UAS-eGFP-sec15/TM3 was obtained from the group of J.A. Knoblich (Institute of Molecular Biotechnology of the Austrian Academy of Sciences). UAS-GalT-GFP was obtained from the group of S. Luschnig (WMU Munster) (Armbruster and Luschnig, 2012).

Fly lines expressing mCherry tagged *ArfGAP1* and *ArfGAP1*^*R50Q*^ under the UAS promoter and the mutated allele of ArfGAP1 (*ArfGAP1*^*5*^) are available upon request (the recipient must pay for shipping) to the Lead Contact, Gregory Emery.

### Cloning

ArfGAP1 was cloned from a cDNA clone obtained from the Berkeley Drosophila Genome Project (RE63354). 5’-GGGGACAAGTTTGTACAAAAAAGCAGGCTTCACCATGGCGAGTCCCAGAACGCG-3’ (forward primer) and 5’-GGGGACCACTTTGTACAAGAAAGCTGGGTCGTTCATCAGCAGATTCCAGG-3’ (reverse primer) were used to amplify the sequence, which was inserted in a Gateway vector pDONR221. ArfGAP1 sequence was later inserted in a destination vector (pDest29) to obtain ArfGAP1-mCherry under the control of the UAS promoter. ArfGAP1^R50Q^-mCherry was generated by Quickchange mutagenesis using 5’-TGCACGCCCAGACTCTGATGCTTGCCGGAG-3’ and 5’-CTCCGGCAAGCATCAGAGTCTGGGCGTGCA-3’ primers.

### Immunofluorescence

Ovaries were dissected in PBS (Phosphate Buffered Saline) and fixed in 200μL of paraformaldehyde 4% in PBS for 20min. After 3 washes in 200μL of Triton X-100 0,3% in PBS, ovaries were incubated in 200μL of bovine serum albumin (BSA) 2% + Triton X-100 0.3% in PBS for 1-3h under agitation at room temperature. Then, ovaries were incubated overnight at 4°C under agitation with the primary antibody (anti-pTyr (4G10) (05-321, Millipore Sigma, 1:40), anti-pERM (3141S, Cell Signaling, 1:100), anti-Fasciclin III (7G10, DSHB, 1:100), anti-DE cadherin (DCAD2, DSHB, 1:50), anti-Bazooka (gift from T.Harris lab, University of Toronto, 1:1500) or anti-Discs Large (4F3,DSHB, 1:100) diluted in BSA 2% + Triton 0.3% in PBS. The next day, after 2 short washes and 2 20min washes under agitation with Triton 0.3% in PBS, ovaries were incubated during 1-3h under agitation at room temperature with secondary antibody (anti-mouse (A11029 or A21424, Invitrogen, 1:500), anti-rabbit (A11008 or A11012, Invitrogen, 1:250) or anti-rat (A11006, Invitrogen, 1:400), Phalloidin (A22287, Invitrogen, 1:1000) and DAPI (D8417-10MG, Sigma, 1:10 000) diluted in BSA 2% + Triton 0.3% in PBS. Then, ovaries were washed again 2 times fast and 2 times during 20min under agitation before mounting on slides with Vectashield (H-1000, Vector Laboratories).

The protocol for staining with LysoTracker RED DND-99 (L7528, Invitrogen) was adapted from DeVorkin and Gorski (2016). Ovaries were dissected in PBS 1X and incubated in 200µL of 50μM LysoTracker RED for 3min in the dark. Then, ovaries were washed 3 times 5min in 200µL of PBS 1X and fixed in 200μL of PFA 4% during 20min protected from light and subsequent steps were performed as previously described.

### Image acquisition and quantitative analysis

Images were acquired using a laser scanning confocal microscope LSM 700 (Carl Zeiss) or a Leica TCS SP8 (Leica Microsystems).

pTyr stainings were quantified as follows: mean pTyr fluorescence intensities were quantified at the membrane at the front of border cell (BC) cluster and at the membrane of nurse cells in three consecutive frames from a z-scan separated by 1μm using original images and the ImageJ software (National Institutes of Health). Lifeact-GFP or Phalloidin staining were used to determine the periphery of the cluster. The ratio between signal at the border cell membranes and nurse cell membranes was calculated. Number of pTyr positive puncta were quantified by manually counting in three consecutive frames from a z-scan separated by 1μm. GFP-Rab7 mean intensity was measured on the entire cluster in three consecutive frames from a z-scan separated by 1μm. For all these analyses, we used the z-scan containing the two polar cells as a central plan. The background, determined outside the egg chamber, was systematically subtracted from the fluorescence intensities determined. Only images with a signal in the linear range were considered for quantification. GFP-Rab7, LysoTracker RED and Sec15 vesicles were analyzed using the Spot plugin on Imaris (Bitplane), on an entire z-scan recorded with frames separated by 1μm. GalT-GFP structures were quantified by manually setting a threshold and automatic counting using the Analyze particles plugin in ImageJ, on a z-projection of five consecutive focal planes separated by 0.5 μm.

### Live imaging

Dissection and primary culture of ovaries expressing LifeAct-GFP were performed as previously described (Prasad et al., 2007). Egg chambers were isolated and cultured in Schneider’s medium containing 200μg/ml of insulin and 15% FBS for live imaging. A z-scan of 3 images separated by 3µm was acquired every 2min over the course of 4 to 6 hours using a spinning disk confocal microscope (Zeiss). For rendering, they were processed on the ImageJ software using the “Maximum Intensity Z*-*projection*”* and the “Contrast and Brightness” function. Drift due to egg chamber movement was corrected on the time-lapse recording of one control border cell cluster using the “TurboReg” plugin in ImageJ. Manual cell tracking and x/y position recording of clusters were performed by using the “Manual Tracking” plugin in ImageJ. Directionality, accumulated distance, Euclidean distance, and velocity were calculated using the chemotaxis tool from Ibidi (http://ibidi.com/software/chemotaxis_and_migration_tool/).

### Migration Quantification

The migration index (M.I.) represents the relative distance, compared to the end of the migration path, migrated by BCs in the egg chamber at stage 10, whereas the completion index (C.I.) represents the ratio of clusters having completed migration, and both were calculated as previously described (Assaker et al., 2010): The Migration Index (M.I.) was calculated with the following formula: (0*n(0%)+0,25*n(25%)+0,5*n(50%)+0,75*n(75%)+1*n(100%))/n(total).

N(100%) corresponds to the number of egg chambers where the cluster reached the oocyte, n(75%), the number of chambers where the cluster migrated to 75% of the final distance, etc. and n(total), the total number of egg chambers. The Completion Index (C.I.) corresponds to the number of egg chambers where the migration was completed divided by the total number of egg chambers: n(100%)/n(total).

### Statistical analysis

Statistical comparisons of means were made by comparing each condition to the adequate control, using the unpaired non-parametric Mann-Whitney test or the Kruskal-Wallis test with Dunn’s correction for multiple comparisons using GraphPad Prism software. For Migration Indexes (M.I.) and Completion Indexes (C.I.), statistical comparisons of means were made by comparing each condition to the adequate control, using the unpaired One-way ANOVA test with correction for multiple comparisons using GraphPad Prism software. P<0.05 is represented by one star (^*^), p<0.01 by two stars (^**^), p<0.001 by three stars (^***^) and p<0.0001 by four stars (^****^). Mean values are quoted ± SEM in figures.

## ACKNOWLEDGMENTS

We thank the Bloomington Stock Collection and the Vienna Drosophila RNAi Center for fly stocks. We thank M. Gonzalez-Gaitan, J.A. Knoblich, S. Luschnig and T. Harris for their generosity in sharing reagents. We thank C. Charbonneau for technical assistance and the entire Emery lab for helpful discussions. This work was supported by grants from the Canadian Institute for Health Research (CIHR; PJT - 175093) and the Natural Sciences and Engineering Research Council of Canada to G.E.. A.B. held a doctoral scholarship from Institute for Research in Immunology and Cancer and from Molecular Biology Program of the University of Montreal. C.Z. held a doctoral scholarship from Fonds de Recherche du Québec – Santé (FRQS).

## FIGURE LEGENDS

**Figure S1:**ArfGAP1 depletion does not affect Golgi integrity, DE-Cadherin, polarity protein nor phosphorylated Moesin. (A) Representative images of border cells expressing GalT-GFP in control condition and after ArfGAP1 depletion. Scale bar: 5µm. (B) Area covered by GalT-positive vesicles in control and ArfGAP1 depleted clusters. Error bars are SEM (6≤n≤18 clusters; N=3 experiments; n.s.: not significant (p>0.05), Student’s t-test). (C) Number of GalT-positive vesicles in control and ArfGAP1 depleted clusters. Error bars are SEM (6≤n≤18 clusters; N=3 experiments; n.s.: not significant (p>0.05), Student’s t-test). (D) Representative images of border cells expressing Lifeact-GFP and labeled with the anti-pMoesin antibody in control condition and after ArfGAP1 depletion. Scale bar: 5µm. (E) Representative images of border cells expressing Lifeact-GFP and labeled with the anti-Discs large antibody in control condition and after ArfGAP1 depletion. Scale bar: 5µm. (F) Representative images of border cells expressing Lifeact-GFP and labeled with the anti-Bazooka antibody in control condition and after ArfGAP1 depletion. Scale bar: 5µm. (G) Representative images of border cells in control and after ArfGAP1 depletion, labeled with the anti-DE-cadherin antibody. Scale bar: 5µm. (H) Representative images of border cells expressing Lifeact-GFP and labeled with the anti-Fasciclin antibody in control condition and after ArfGAP1 depletion. Scale bar: 5µm. (I) Number of Sec15-positive vesicles in control and ArfGAP1 depleted clusters. Error bars are SEM (N=3 experiments; n.s.: not significant (p>0.05), Mann-Whitney test).

**Movie S1:** Time-Lapse recording of the migration of a cluster expressing Lifeact-GFP (related to Figure 1). Frames were acquired every 2min with a 20x objective.

**Movie S2:** Time-Lapse recording of an ArfGAP1 depleted cluster expressing Lifeact-GFP (related to Figure 1). The cluster can be seen extending protrusions in different directions and tumbling but does not migrate. Frames were acquired every 2min with a 20x objective.

**Movie S3:** Time-Lapse recording of an ArfGAP1 depleted cluster expressing Lifeact-GFP (related to Figure 1). The cluster starts migrating but goes in different direction and not toward the oocyte. Frames were acquired every 2min with a 20x objective.

## REFERENCES

Adarska, P., L. Wong-Dilworth, and F. Bottanelli. 2021. ARF GTPases and Their Ubiquitous Role in Intracellular Trafficking Beyond the Golgi. Front Cell Dev Biol. 9:679046.

Armbruster, K., and S. Luschnig. 2012. The Drosophila Sec7 domain guanine nucleotide exchange factor protein Gartenzwerg localizes at the cis-Golgi and is essential for epithelial tube expansion. J Cell Sci. 125:1318–1328.

Assaker, G., D. Ramel, S.K. Wculek, M. Gonzalez-Gaitan, and G. Emery. 2010. Spatial restriction of receptor tyrosine kinase activity through a polarized endocytic cycle controls border cell migration. Proc Natl Acad Sci U S A. 107:22558–22563.

Bianco, A., M. Poukkula, A. Cliffe, J. Mathieu, C.M. Luque, T.A. Fulga, and P. Rorth. 2007. Two distinct modes of guidance signalling during collective migration of border cells. Nature. 448:362–365.

Cai, D., S.C. Chen, M. Prasad, L. He, X. Wang, V. Choesmel-Cadamuro, J.K. Sawyer, G. Danuser, and D.J. Montell. 2014. Mechanical feedback through E-cadherin promotes direction sensing during collective cell migration. Cell. 157:1146–1159.

Caldieri, G., M.G. Malabarba, P.P. Di Fiore, and S. Sigismund. 2018. EGFR Trafficking in Physiology and Cancer. Prog Mol Subcell Biol. 57:235–272.

Capuana, L., A. Bostrom, and S. Etienne-Manneville. 2020. Multicellular scale front-to-rear polarity in collective migration. Curr Opin Cell Biol. 62:114–122.

Civiero, L., and L. Bubacco. 2012. Human leucine-rich repeat kinase 1 and 2: intersecting or unrelated functions? Biochem Soc Trans. 40:1095–1101.

Cobreros-Reguera, L., A. Fernandez-Minan, C.H. Fernandez-Espartero, H. Lopez-Schier, A. Gonzalez-Reyes, and M.D. Martin-Bermudo. 2010. The Ste20 kinase misshapen is essential for the invasive behaviour of ovarian epithelial cells in Drosophila. EMBO Rep. 11:943–949.

Cukierman, E., I. Huber, M. Rotman, and D. Cassel. 1995. The ARF1 GTPase-activating protein: zinc finger motif and Golgi complex localization. Science. 270:1999–2002.

Dodson, M.W., T. Zhang, C. Jiang, S. Chen, and M. Guo. 2012. Roles of the Drosophila LRRK2 homolog in Rab7-dependent lysosomal positioning. Hum Mol Genet. 21:1350–1363.

Duchek, P., and P. Rorth. 2001. Guidance of cell migration by EGF receptor signaling during Drosophila oogenesis. Science. 291:131–133.

Duchek, P., K. Somogyi, G. Jekely, S. Beccari, and P. Rorth. 2001. Guidance of cell migration by the Drosophila PDGF/VEGF receptor. Cell. 107:17–26.

Fernandez-Espartero, C.H., D. Ramel, M. Farago, M. Malartre, C.M. Luque, S. Limanovich, S. Katzav, G. Emery, and M.D. Martin-Bermudo. 2013. GTP exchange factor Vav regulates guided cell migration by coupling guidance receptor signalling to local Rac activation. J Cell Sci. 126:2285–2293.

Gomez-Suaga, P., P. Rivero-Rios, E. Fdez, M. Blanca Ramirez, I. Ferrer, A. Aiastui, A. Lopez De Munain, and S. Hilfiker. 2014. LRRK2 delays degradative receptor trafficking by impeding late endosomal budding through decreasing Rab7 activity. Hum Mol Genet. 23:6779–6796.

Haeger, A., K. Wolf, M.M. Zegers, and P. Friedl. 2015. Collective cell migration: guidance principles and hierarchies. Trends Cell Biol. 25:556–566.

Han, D.D., D. Stein, and L.M. Stevens. 2000. Investigating the function of follicular subpopulations during Drosophila oogenesis through hormone-dependent enhancer-targeted cell ablation. Development. 127:573–583.

Hanafusa, H., K. Ishikawa, S. Kedashiro, T. Saigo, S. Iemura, T. Natsume, M. Komada, H. Shibuya, A. Nara, and K. Matsumoto. 2011. Leucine-rich repeat kinase LRRK1 regulates endosomal trafficking of the EGF receptor. Nat Commun. 2:158.

Hanafusa, H., T. Yagi, H. Ikeda, N. Hisamoto, T. Nishioka, K. Kaibuchi, K. Shirakabe, and K. Matsumoto. 2019. LRRK1 phosphorylation of Rab7 at S72 links trafficking of EGFR-containing endosomes to its effector RILP. J Cell Sci. 132.

Heasman, S.J., and A.J. Ridley. 2008. Mammalian Rho GTPases: new insights into their functions from in vivo studies. Nat Rev Mol Cell Biol. 9:690–701.

Ishikawa, K., A. Nara, K. Matsumoto, and H. Hanafusa. 2012. EGFR-dependent phosphorylation of leucinerich repeat kinase LRRK1 is important for proper endosomal trafficking of EGFR. Mol Biol Cell. 23:1294–1306.

Jackson, C.L., and S. Bouvet. 2014. Arfs at a glance. J Cell Sci. 127:4103–4109.

Jafar-Nejad, H., H.K. Andrews, M. Acar, V. Bayat, F. Wirtz-Peitz, S.Q. Mehta, J.A. Knoblich, and H.J. Bellen. 2005. Sec15, a component of the exocyst, promotes notch signaling during the asymmetric division of Drosophila sensory organ precursors. Dev Cell. 9:351–363.

Janssens, K., H.H. Sung, and P. Rorth. 2010. Direct detection of guidance receptor activity during border cell migration. Proc Natl Acad Sci U S A. 107:7323–7328.

Jekely, G., H.H. Sung, C.M. Luque, and P. Rorth. 2005. Regulators of endocytosis maintain localized receptor tyrosine kinase signaling in guided migration. Dev Cell. 9:197–207.

Khadilkar, R.J., D. Rodrigues, R.D. Mote, A.R. Sinha, V. Kulkarni, S.S. Magadi, and M.S. Inamdar. 2014. ARF1-GTP regulates Asrij to provide endocytic control of Drosophila blood cell homeostasis. Proc Natl Acad Sci U S A. 111:4898–4903.

Laflamme, C., G. Assaker, D. Ramel, J.F. Dorn, D. She, P.S. Maddox, and G. Emery. 2012. Evi5 promotes collective cell migration through its Rab-GAP activity. J Cell Biol. 198:57–67.

Lee, S.Y., J.S. Yang, W. Hong, R.T. Premont, and V.W. Hsu. 2005. ARFGAP1 plays a central role in coupling COPI cargo sorting with vesicle formation. J Cell Biol. 168:281–290.

Li, Q., L. Shen, T. Xin, W. Xiang, W. Chen, Y. Gao, M. Zhu, L. Yu, and M. Li. 2009. Role of Scrib and Dlg in anterior-posterior patterning of the follicular epithelium during Drosophila oogenesis. BMC Dev Biol. 9:60.

Liu, W., R. Duden, R.D. Phair, and J. Lippincott-Schwartz. 2005. ArfGAP1 dynamics and its role in COPI coat assembly on Golgi membranes of living cells. J Cell Biol. 168:1053–1063.

Malik, A.U., A. Karapetsas, R.S. Nirujogi, S. Mathea, D. Chatterjee, P. Pal, P. Lis, M. Taylor, E. Purlyte, R. Gourlay, M. Dorward, S. Weidlich, R. Toth, N.K. Polinski, S. Knapp, F. Tonelli, and D.R. Alessi. 2021. Deciphering the LRRK code: LRRK1 and LRRK2 phosphorylate distinct Rab proteins and are regulated by diverse mechanisms. Biochem J. 478:553–578.

McDonald, J.A., E.M. Pinheiro, L. Kadlec, T. Schupbach, and D.J. Montell. 2006. Multiple EGFR ligands participate in guiding migrating border cells. Dev Biol. 296:94–103.

McDonald, J.A., E.M. Pinheiro, and D.J. Montell. 2003. PVF1, a PDGF/VEGF homolog, is sufficient to guide border cells and interacts genetically with Taiman. Development. 130:3469–3478.

Mishra, A.K., J.A. Mondo, J.P. Campanale, and D.J. Montell. 2019. Coordination of protrusion dynamics within and between collectively migrating border cells by myosin II. Mol Biol Cell. 30:2490–2502.

Montell, D.J. 2003. Border-cell migration: the race is on. Nat Rev Mol Cell Biol. 4:13–24.

Montell, D.J., W.H. Yoon, and M. Starz-Gaiano. 2012. Group choreography: mechanisms orchestrating the collective movement of border cells. Nat Rev Mol Cell Biol. 13:631–645.

Niewiadomska, P., D. Godt, and U. Tepass. 1999. DE-Cadherin is required for intercellular motility during Drosophila oogenesis. J Cell Biol. 144:533–547.

Peercy, B.E., and M. Starz-Gaiano. 2020. Clustered cell migration: Modeling the model system of Drosophila border cells. Semin Cell Dev Biol. 100:167–176.

Pinheiro, E.M., and D.J. Montell. 2004. Requirement for Par-6 and Bazooka in Drosophila border cell migration. Development. 131:5243–5251.

Plutoni, C., S. Keil, C. Zeledon, L.E.A. Delsin, B. Decelle, P.P. Roux, S. Carreno, and G. Emery. 2019. Misshapen coordinates protrusion restriction and actomyosin contractility during collective cell migration. Nat Commun. 10:3940.

Prasad, M., and D.J. Montell. 2007. Cellular and molecular mechanisms of border cell migration analyzed using time-lapse live-cell imaging. Dev Cell. 12:997–1005.

Raghu, P., E. Coessens, M. Manifava, P. Georgiev, T. Pettitt, E. Wood, I. Garcia-Murillas, H. Okkenhaug, D. Trivedi, Q. Zhang, A. Razzaq, O. Zaid, M. Wakelam, C.J. O’Kane, and N. Ktistakis. 2009. Rhabdomere biogenesis in Drosophila photoreceptors is acutely sensitive to phosphatidic acid levels. J Cell Biol. 185:129–145.

Ramel, D., X. Wang, C. Laflamme, D.J. Montell, and G. Emery. 2013. Rab11 regulates cell-cell communication during collective cell movements. Nature cell biology. 15:317–324.

Roberto, G.M., and G. Emery. 2022. Directing with restraint: Mechanisms of protrusion restriction in collective cell migrations. Semin Cell Dev Biol. 129:75–81.

Rodrigues, F.F., and T.J.C. Harris. 2019. Key roles of Arf small G proteins and biosynthetic trafficking for animal development. Small GTPases. 10:403–410.

Rodriguez-Fernandez, I.A., and E.C. Dell’Angelica. 2015. Identification of Atg2 and ArfGAP1 as Candidate Genetic Modifiers of the Eye Pigmentation Phenotype of Adaptor Protein-3 (AP-3) Mutants in Drosophila melanogaster. PLoS One. 10:e0143026.

SenGupta, S., C.A. Parent, and J.E. Bear. 2021. The principles of directed cell migration. Nat Rev Mol Cell Biol. 22:529–547.

Stafa, K., A. Trancikova, P.J. Webber, L. Glauser, A.B. West, and D.J. Moore. 2012. GTPase activity and neuronal toxicity of Parkinson’s disease-associated LRRK2 is regulated by ArfGAP1. PLoS Genet. 8:e1002526.

Usmani, A., F. Shavarebi, and A. Hiniker. 2021. The Cell Biology of LRRK2 in Parkinson’s Disease. Mol Cell Biol. 41.

Wan, P., D. Wang, J. Luo, D. Chu, H. Wang, L. Zhang, and J. Chen. 2013. Guidance receptor promotes the asymmetric distribution of exocyst and recycling endosome during collective cell migration. Development. 140:4797–4806.

Wang, H., X. Guo, X. Wang, X. Wang, and J. Chen. 2020. Supracellular Actomyosin Mediates Cell-Cell Communication and Shapes Collective Migratory Morphology. iScience. 23:101204.

Wang, H., Z. Qiu, Z. Xu, S.J. Chen, J. Luo, X. Wang, and J. Chen. 2018. aPKC is a key polarity determinant in coordinating the function of three distinct cell polarities during collective migration. Development. 145.

Wucherpfennig, T., M. Wilsch-Brauninger, and M. Gonzalez-Gaitan. 2003. Role of Drosophila Rab5 during endosomal trafficking at the synapse and evoked neurotransmitter release. J Cell Biol. 161:609–624.

Xiong, Y., C. Yuan, R. Chen, T.M. Dawson, and V.L. Dawson. 2012. ArfGAP1 is a GTPase activating protein for LRRK2: reciprocal regulation of ArfGAP1 by LRRK2. J Neurosci. 32:3877–3886.

Zeledon, C., X. Sun, C. Plutoni, and G. Emery. 2019. The ArfGAP Drongo Promotes Actomyosin Contractility during Collective Cell Migration by Releasing Myosin Phosphatase from the Trailing Edge. Cell Rep. 28:3238–3248 e3233.

Zirin, J., Y. Hu, L. Liu, D. Yang-Zhou, R. Colbeth, D. Yan, B. Ewen-Campen, R. Tao, E. Vogt, S. VanNest, C. Cavers, C. Villalta, A. Comjean, J. Sun, X. Wang, Y. Jia, R. Zhu, P. Peng, J. Yu, D. Shen, Y. Qiu, L. Ayisi, H. Ragoowansi, E. Fenton, S. Efrem, A. Parks, K. Saito, S. Kondo, L. Perkins, S.E. Mohr, J. Ni, and N. Perrimon. 2020. Large-Scale Transgenic Drosophila Resource Collections for Loss-and Gain-of-Function Studies. Genetics. 214:755–767.

