## Supplementary figures and images for "ArfGAP1 regulates the endosomal sorting of guidance receptors to promote directed collective cell migration in vivo"

### Figure S1

#### Supplementary Figure

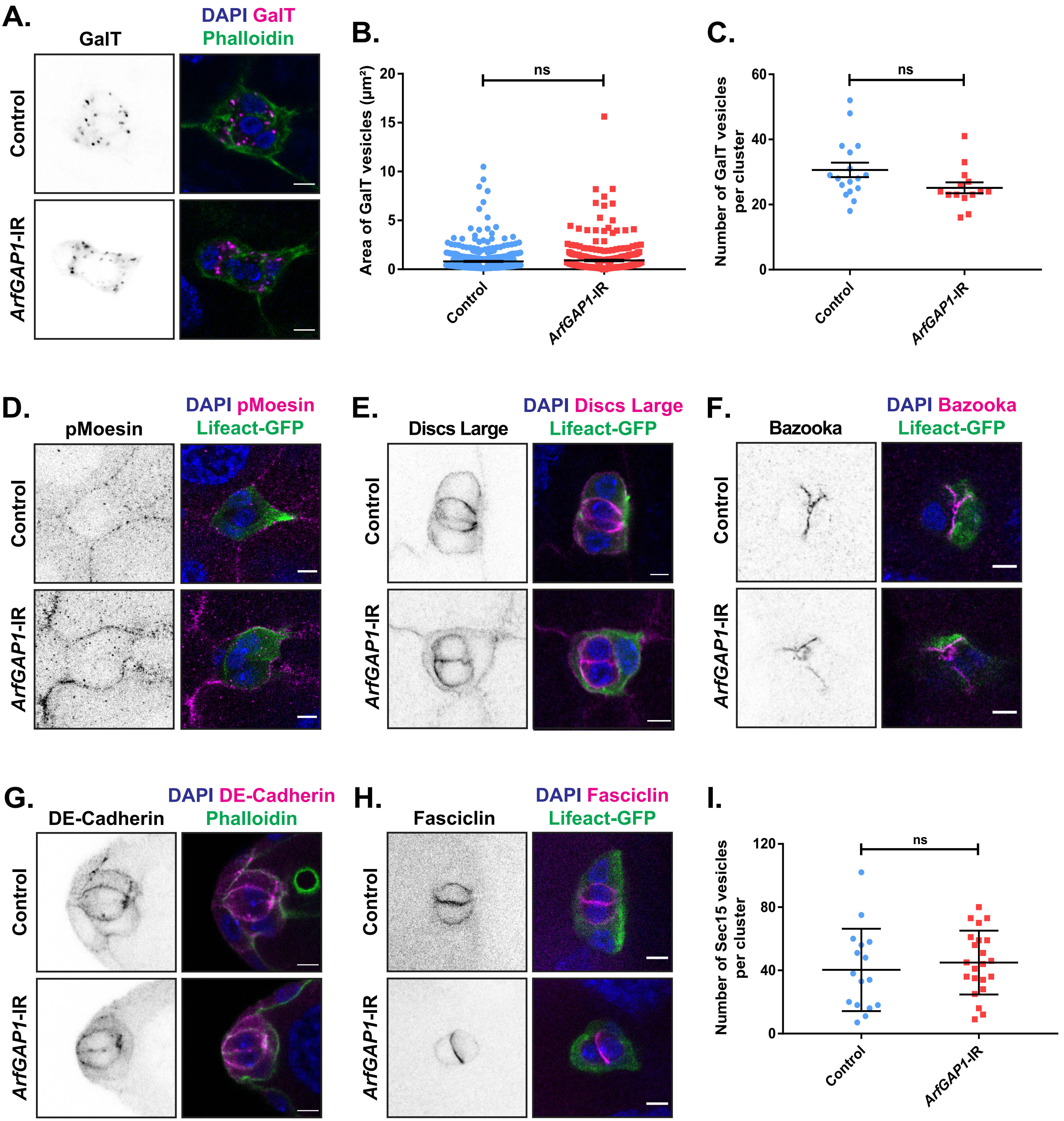
